# An *in vitro* assay to investigate venom neurotoxin activity on muscle-type nicotinic acetylcholine receptor activation and for the discovery of toxin-inhibitory molecules

**DOI:** 10.1101/2023.04.28.538762

**Authors:** Rohit N Patel, Rachel H Clare, Line Ledsgaard, Mieke Nys, Jeroen Kool, Andreas H Laustsen, Chris Ulens, Nicholas R Casewell

## Abstract

Snakebite envenoming is a neglected tropical disease that causes over 100,000 deaths annually. Envenomings result in variable pathologies, but systemic neurotoxicity is among the most serious and is currently only treated with difficult to access and variably efficacious commercial antivenoms. Venom-induced neurotoxicity is often caused by α-neurotoxins antagonising the muscle-type nicotinic acetylcholine receptor (nAChR), a ligand-gated ion channel. Discovery of therapeutics targeting α-neurotoxins is hampered by relying on binding assays that do not reveal restoration of receptor activity or more costly and/or lower throughput electrophysiology-based approaches. Here, we report the validation of a screening assay for nAChR activation using immortalised TE671 cells expressing the γ-subunit containing muscle-type nAChR and a fluorescent dye that reports changes in cell membrane potential. Assay validation using traditional nAChR agonists and antagonists, which either activate or block ion fluxes, was consistent with previous studies. We then characterised antagonism of the nAChR by a variety of elapid snake venoms that cause muscle paralysis in snakebite victims, before defining the toxin-inhibiting activities of commercial antivenoms, and new types of snakebite therapeutic candidates, namely monoclonal antibodies, decoy receptors, and small molecules. Our findings show robust evidence of assay uniformity across 96-well plates and highlight the amenability of this approach for the future discovery of new snakebite therapeutics via screening campaigns. The described assay therefore represents a useful first-step approach for identifying α-neurotoxins and their inhibitors in the context of snakebite envenoming, and it should provide wider value for studying modulators of nAChR activity from other sources.

## 1. Introduction

Snakebite envenoming is a neglected tropical disease that is responsible for causing over 100,000 deaths and 400,000 disabilities each year [1]. To achieve the targets set out in the World Health Organization’s (WHO’s) snakebite roadmap to halve deaths and disability by 2030, more effective, affordable, and accessible treatments are urgently needed [2]. Howev, snake venom variation acts as a barrier to the development of broadly effective therapeutics because inter-specific toxin variation results in a diversity of pathogenic drug targets that cause variable envenoming pathologies in snakebite victims, *i.e.*, haemotoxicity, cytotoxicity, and/or neurotoxicity [3]. Snake venom composition is dictated by variable representation by several toxin families, such as snake venom metalloproteinases (SVMPs), snake venom serine proteases (SVSPs), phospholipases A_2_ (PLA_2_s), and three-finger toxins (3FTxs) [4]. The latter two are usually of greatest significance in medically important elapid snake venoms [5], with highly abundant 3FTx isoforms often responsible for causing potentially lethal systemic neurotoxicity [6].

3FTxs are broadly subdivided by their structure and site of action into different subcategories. 3FTxs that exert their activity by binding to nicotinic acetylcholine receptors (nAChRs) located on the post-synaptic membranes of neuromuscular junctions are collectively known as α-neurotoxins (α-NTxs) [7]. α-NTxs are further subdivided based on their structure into long-chain (Lc-α-NTx), short-chain (Sc-α-NTx), non-conventional, and weak α-NTxs [6]. nAChRs are pentameric ligand-gated ion channels gated by the binding of the neurotransmitter acetylcholine (ACh) [8]. The nAChR located at the neuromuscular junction (referred to as ‘muscle-type’) consists of a combination of two α1 subunits with β1, δ, and either a γ subunit during foetal development (foetal) or a ε subunit thereafter (adult) [9]. Muscle-type nAChR activation results in skeletal muscle contraction, while binding of α-NTxs, which bind with high affinity and can have lengthy dissociation times [10], prevents activation by blocking Ach binding, resulting in neurotoxicity, which presents clinically in snakebite victims as ptosis, muscular paralysis, and respiratory depression [11,12].

Commercially available antivenoms are the only approved specific treatment for snakebite envenoming. They consist of polyclonal antibodies purified from the plasma/sera of animals immunised with sub-toxic doses of venom [13] and have proven to be effective at preventing life-threatening signs of systemic envenoming if delivered promptly [14]. However, current antivenoms have several limitations associated with them, including poor dose efficacy, limited cross-snake species efficacy, high frequency of adverse reactions due to their heterologous nature, and low affordability and accessibility to tropical snakebite victims [15,16].

In recent years, several new approaches to either improve, supplement, or replace existing antivenoms have been described [17–21]. Because neurotoxic envenoming can rapidly become life-threatening, toxins that act on the nAChR are priority targets for the discovery of novel therapeutics. Investigation of snake toxin action on nAChR functioning is traditionally carried out using electrophysiological recordings [22] and/or recordings from *ex vivo* nerve-muscle preparations [23]. However, these techniques are laborious, low-throughput, and resource-intensive, and are therefore barriers to identifying novel neurotoxin-inhibiting molecules (*e.g.*, monoclonal antibodies, peptides, and/or small molecule drugs). More recently, automated patch-clamping has been introduced as a high-throughput method that allows for similar types of electrophysiological recordings [24–26]. However, this approach requires sophisticated equipment that is not available in most laboratories. Alternative methods to investigate toxin-nAChR interactions have been developed, including the use of mimotopes of the α1 nAChR subunit toxin binding site [27], a binding assay using purified nAChR from the electric organ of *Torpedo* species [28], and the use of acetylcholine binding protein (AChBP), a soluble protein from mollusc species, as a proxy for the nAChR [29]. However, each of these alternative approaches examines receptor binding rather than functionality.

A promising approach using immortalised TE671 cells expressing the foetal muscle-type nAChR [30] and a membrane potential dye to report receptor activation [31] has been used to investigate the activity of a Lc-α-NTx isolated from black mamba (*Dendroaspis polylepis*) venom [32]. The membrane potential dye moves intracellularly due to cation influx after receptor activation and binds to intracellular proteins and lipids resulting in an increase in fluorescence. This allows measurements of nAChR activation using an affordable plate reader and without the need for specialised electrophysiology equipment or facilities. TE671 cells have been widely used to investigate muscle-type nAChR function using patch-clamp electrophysiology [33–35] and, with membrane potential dye, have been used to investigate the nAChR activity of natural compounds [36–38], and to identify neuronal nAChR antagonists of relevance for tobacco addiction [39]. In this study, we exploited the assay potential of TE671 cells incubated with a membrane potential dye and validated this approach as a tool for: i) characterising the nAChR antagonism of crude snake venoms and isolated snake venom toxins, and ii) use as a 96-well plate *in vitro* assay platform for the discovery of novel toxin-inhibiting therapeutics (Fig. 1).

**Fig. 1.**
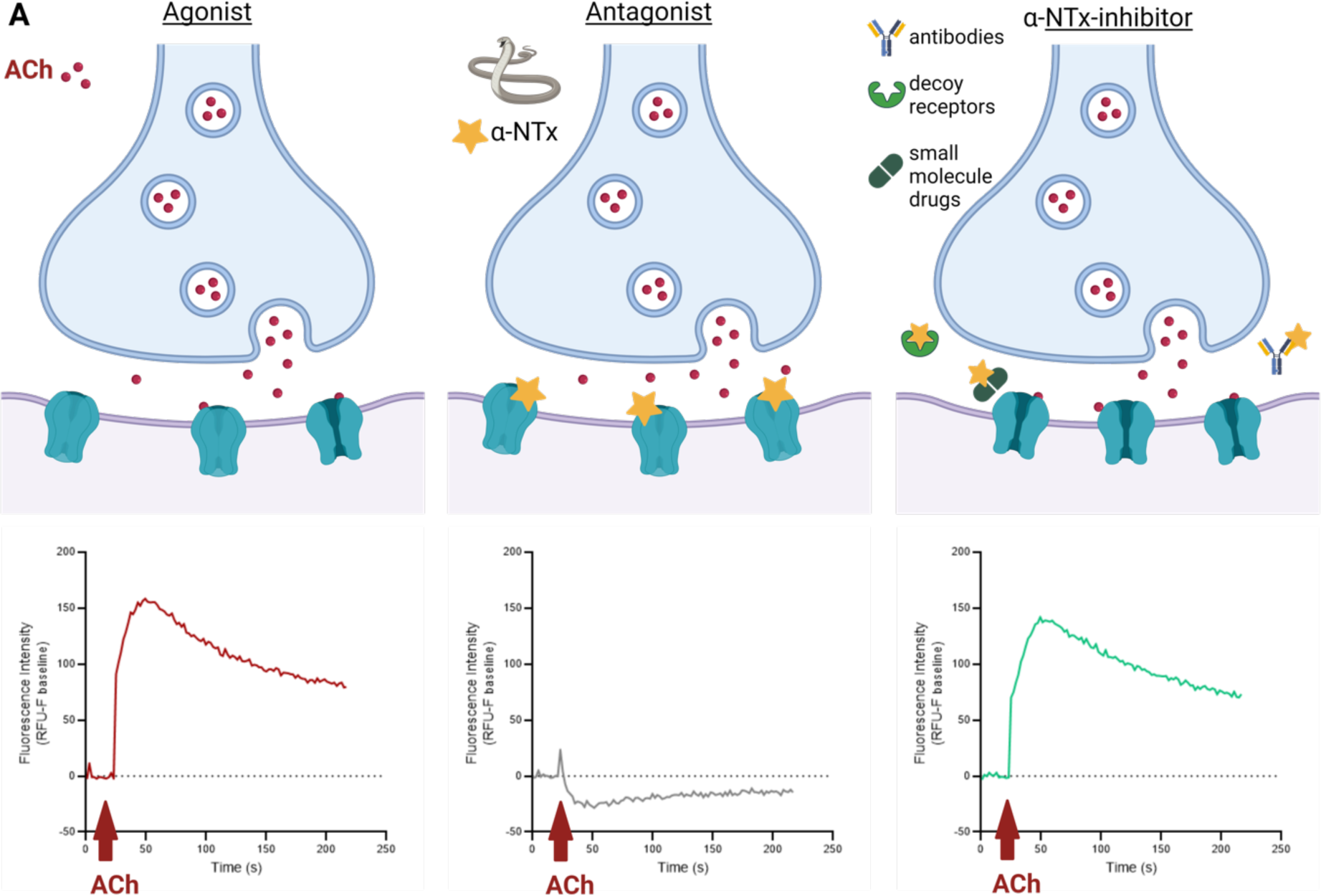
Schematic of the neuromuscular junction and overview of corresponding readouts from the developed assay of nAChR activation. The schematics of the neuromuscular junction (top panels) demonstrate the release of ACh from the pre-synaptic neuron and binding of ACh to the post-synaptic membrane in the absence of α-NTxs (left, Agonist), in the presence of α-NTxs (middle, Antagonist), and in the presence of both α-NTxs and α-NTx-inhibitors (right, α-NTx-inhibitor). Underneath each illustration is the typical fluorescent response using the assay of a well in a 96-well plate containing TE671 cells when ACh alone (Agonist), ACh and α-NTx (Antagonist) and ACh, α-NTx and α-NTx-inhibitors (α-NTx-inhibitors: antibodies, decoy receptors, and small molecules) are added. All schematics were created with BioRender.

### 1.2. Experimental procedures

#### 2.1 Materials

##### 2.1.1 Venoms

Crude venoms were extracted from adult wild-caught specimens maintained in the herpetarium facility of the Centre for Snakebite Research & Interventions at the Liverpool School of Tropical Medicine (LSTM) (Liverpool, UK). The facility and its protocols for the husbandry of snakes are approved and inspected by the UK Home Office and the LSTM and University of Liverpool Animal Welfare and Ethical Review Boards. Venoms of the following elapid snake species listed with their common name and country of origin were used: *Dendroaspis polylepis* (black mamba, Tanzania), *Dendroaspis viridis* (Western green mamba, Togo), *Dendroaspis angusticeps* (Eastern green mamba, Tanzania), *Dendroaspis jamesoni jamesoni* (Jameson’s mamba, western subspecies, Cameroon), *Dendroaspis jamesoni kaimosae* (Jameson’s mamba, eastern subspecies, Uganda), *Naja haje* (Egyptian cobra, Uganda), *Naja subfulva* (brown forest cobra, Uganda), and *Naja nivea* (cape cobra, South Africa). After extraction, venoms were immediately stored at -20 °C, lyophilised overnight, and stored long-term at 4 °C. Subsequent lyophilised extractions from each specimen were pooled with previous extractions. Concentrated stock solutions were created by reconstituting the lyophilised powder in PBS (10010023, Gibco, Thermo Fisher Scientific, Paisley, UK) and stored at -80 °C. Concentrations of venoms used in all experiments are expressed as the dry mass of lyophilised venom per mL of diluent.

##### 2.1.2 nAChR agonists and antagonists

The following nAChR agonists were commercially acquired: acetylcholine chloride (A6625, Sigma-Aldrich, Gillingham, UK), nicotine ditartrate (GSK5294, Sigma-Aldrich, Gillingham, UK), and epibatidine dihydrochloride (AOB5901, Aobius, Gloucester, MA, USA). The Sc-α-NTx ‘SHORT NEUROTOXIN alpha (NP)’ (listed with the recommended name ‘short neurotoxin 1’ (sNTx1) on the UniProt database; P01426) isolated from *Naja pallida* venom was purchased from Latoxan (L8101, Valence, France), and the Lc-α-NTx, ‘α-bungarotoxin’ (α-BgTx), isolated from *Bungarus multicinctus* venom was purchased from Biotium (0010-1, Fremont, CA, USA).

##### 2.1.3 Toxin-inhibiting molecules

The polyclonal antibody-based antivenoms EchiTAbG (batch EOG001740, expiry date October 2018, MicroPharm, Newcastle Emlyn, UK) and SAIMR (South African Institute for Medical Research) Polyvalent Snake antivenom (batch BF00546, expiry date November 2017, South African Vaccine Producers [SAVP], Johannesburg, South Africa) were obtained from the LSTM herpetarium via donation from Public Health England (London, UK). AChBP from *Lymnaea stagnalis* (*Ls*-AchBP) was prepared as previously described [17], as were the fully human monoclonal antibodies (mAbs) 2551_01_A12, 2554_01_D11 and 367_01_H01 in IgG1 format [21]. Samples of the various small molecule drugs used for screening were obtained by request from the Open Chemical Repository of the Developmental Therapeutics Program (https://dtp.cancer.gov) (Division of Cancer Treatment and Diagnosis, National Cancer Institute, Rockville, MD, USA), except for nicotine (see section 2.1.2) and varespladib (SML1100, Sigma-Aldrich, Gillingham, UK). These were selected based on their implied potential as α-NTx-inhibitors in previous studies [40–42]. Stock solutions of small molecules were created using dimethyl sulfoxide (DMSO) (D8418, Sigma-Aldrich, Gillingham, UK), and working solutions did not exceed 1% DMSO.

##### 2.1.4 Cell line

The immortalised TE671 cell line (RRID: CVCL_1756) as used in Ngum et al. [43] was gifted by Dr Ian Mellor (University of Nottingham, UK) and originally obtained from the European Collection of Authenticated Cell Cultures (ECACC; catalogue no. 89071904). TE671 is a rhabdomyosarcoma cell line that natively expresses the foetal muscle-type nAChR (γ-subunit containing) [30].

### 2.2 Culture of TE671 cells

All further reagents were acquired from Gibco, Thermo Fisher Scientific, Paisley, UK, unless stated otherwise. TE671 cells were maintained using a culture medium consisting of DMEM (high glucose, with GlutaMAX supplement, 10566016) supplemented with 10% FBS (qualified, Brazil origin, 10270106) and 1% penicillin-streptomycin solution (5000 units/mL penicillin, 5 mg/mL streptomycin, 15070063). Cells were cultured in 75 cm^2^ cell culture flasks (83.3911, Sarstedt, Nümbrecht, Germany) and incubated at 37 °C/5% CO_2_ until ∼90% confluence was reached, upon which cells were dislodged from the flask with 4 mL TrypLE express enzyme (1x, no phenol red, 12604013). The suspension was added to 10 mL culture medium and centrifuged for 5 minutes (min) at 300 × g. The supernatant was removed, and the pellet resuspended in 5 mL culture medium. Cell suspensions of different flasks were pooled, counted using an automated cell counter (Luna II, Logos Biosystems, Villeneuve-d’Ascq, France) and further culture medium added to reach a count of 3×10^4^-4×10^4^ cells/100 µL. Next, 100 µL cell suspension was pipetted to the wells of black walled, clear bottom, tissue culture treated 96-well plates (655090, Greiner Bio One, Stonehouse, UK) and incubated overnight at 37 °C/5% CO_2_.

### 2.3 Membrane potential assay of nAChR activation

The following method was adapted from Fitch et al. [24] and Wang et al. [25] and, as in section 2.2, all reagents were acquired from Gibco, Thermo Fisher Scientific, Paisley, UK, unless stated otherwise. One vial of FLIPR membrane potential dye (Component A, Explorer Kit Blue, R8042, Molecular Devices, San Jose, CA, USA) was dissolved in 36 mL assay buffer to create the dye solution. Assay buffer consisted of 1x HBSS (made from 10x solution [14065049] as per manufacturer’s instruction by diluting with distilled water and addition of NaHCO_3_ [7.5% solution, 25080094] to a final concentration of 4.17 mM) supplemented with 20 mM HEPES (1 M solution, 15630056), 0.5 µM atropine (A0132, Sigma-Aldrich, Gillingham, UK), adjusted to pH 7.1 with 1 M NaOH, and then sterile filtered. Assay buffer was then used to create all further solutions. Culture medium was removed from the cell plate, replaced with 50 µL dye solution and incubated for 30 min at 37 °C/5% CO_2_. When investigating venom/toxin inhibition, the solutions of venom, toxin, toxin-inhibitor, or combinations thereof were concurrently incubated for 30 min at 37 °C/5% CO_2_ prior to addition to the cell plate. Next, 50 µL of control or venom/toxin or venom/toxin + toxin-inhibitor solutions were transferred to each well, and the cell plate further incubated for 15 min at 37 °C/5% CO_2_. The cell plate was then acclimatised for 15 min at room temperature before recording. Next, 60 µL nAChR agonist solution or assay buffer was added to the wells of a clear, v-bottom 96-well plate (651201, Greiner Bio One, Stonehouse, UK) to create a reagent plate and was then added to the appropriate tray, along with the cell plate and a rack of pipette tips (black, 96-well configuration, 9000-0911, Molecular Devices, San Jose, CA, USA), to a FlexStation 3 multi-mode microplate reader (Molecular Devices, San Jose, CA, USA) controlled by SoftMax Pro 7.1 software (Molecular Devices, San Jose, CA, USA). The reader records a column of the 96 well plate for a time set by the user and houses an automated pipetting system that allows the addition of solution from the reagent plate to the cell plate at a set time during the recording. After the column is recorded the adjacent column is then recorded in the same manner. For the purposes of this study this allowed a baseline recording followed by the addition of an agonist solution and the recording of changes in dye fluorescence immediately following this addition. Excitation, cut-off, and emission wavelengths were set at 530, 550, and 565 nm respectively. Recordings of plates were carried out at room temperature using a reading time of 214 seconds (s) and interval time of 2 s to give a total of 108 readings per well with compound transfer (addition of agonist solution) of 50 µL to each well after 20 s baseline recording.

### 2.4 Data and Statistical Analysis

Fluorescent responses for each well were measured by the software in relative fluorescent units (RFUs), and values were determined by calculating the baseline fluorescence (F_baseline_, the mean of the first 20 s of responses) and subtracting this from the maximum fluorescent response (F_max_) for the remainder of the recording for each well (F_max_-F_baseline_). As different wells can have different starting RFU values, this normalisation approach ensured that the responses detected from each well could be compared on the same scale.

For experiments to profile agonists and antagonists, assay buffer alone was included as a control. For all subsequent work, 10 µM ACh was used as the control agonist, and all data points normalised to this agonist alone control. For experiments with isolated α-NTxs, 10 µM ACh was applied after incubation with varying concentrations of toxin. Crude venom experiments were carried out and normalised in the same way. Experiments with toxin-inhibitors included the ACh control (agonist alone), antagonist (venom or toxin) + ACh, and toxin-inhibitor + ACh. The screen of a panel of potential small molecule α-NTx-toxin inhibitors included the above controls, as well as α-BgTx controls of 30 nM (MIN) and 3 nM (MID). For all experiments with toxin-inhibitors, the concentrations of antagonist and ACh were kept consistent and co-incubated with varying concentrations of toxin-inhibitor. The data was then normalised to the mean ACh control (100% signal) and ACh + venom/toxin (0% signal) controls using equation (1) with venom/toxin-inhibitory activity represented by recovery towards the 100% ACh signal.

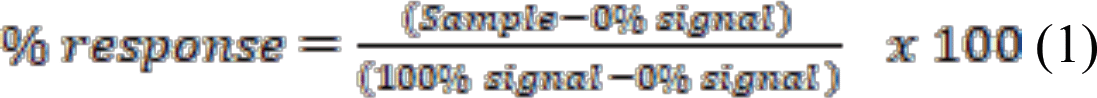

All experiments had 3-8 replicates per plate and were repeated on three separate plates using different cell passage numbers (4-12), and each repeat was carried out on a different day. Normalised data was combined across plates by calculating the mean of the replicates of each plate to give a single value for each plate (n = 1) and subsequently calculating the mean of these combined values. As experiments were repeated three times, all experiments had n = 3, and each data point was plotted as the mean ± SD. Plate uniformity studies were carried out as previously described [44], with assay quality measured using Z prime (Z’) analysis with an acceptance criterion of ≥ 0.4 [45]. Each data point across the three plates was plotted individually (rather than mean ± SD), so the variability of responses across the plate could be visualised. All data analysis, graph plotting, and application of non-linear regression equations (2) and (3) to fit curves were carried out using Prism 9 (GraphPad, San Diego, CA, USA).

The following non-linear regression equation was applied to fit a curve to concentration-response plots to generate EC_50_ values:

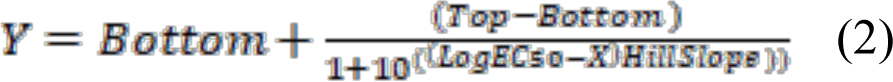

The following non-linear regression equation was applied to fit a curve to concentration-inhibition plots to generate IC_50_ values:

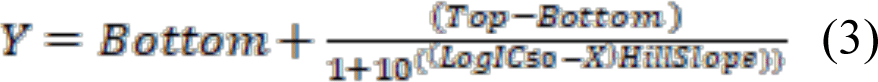

## 3. Results

### 3.1 nAChR agonists produce fluorescent responses in TE671 cells that are blocked by known nAChR antagonists

To determine whether our modifications to the previously described assay protocols produced consistent data, we validated the assay using several known nAChR agonists (ACh, nicotine, and epibatidine) and antagonists (snake venom Lc-α-NTxs and Sc-α-NTxs) of the muscle-type nAChR, alongside assay buffer alone (negative control). The incubation of TE671 cells with this negative control plus the membrane potential dye resulted in a slight decrease in the RFU readings that remained slightly below F_baseline_ levels for the remainder of the recording, indicating that the addition of solution itself causes a small decrease in fluorescence (Fig. 1). The addition of the three nAChR agonists resulted in concentration-dependent increases in fluorescence. The profile of the responses to all agonists typically reached F_max_ approximately 45 s after addition, followed by a slow decay for the remainder of the recording to up to 50% of the peak (Fig. 2A). Concentration-response curves revealed a rank order of potency of epibatidine > ACh > nicotine (Fig. 2B, Table 1). In the case of epibatidine, increasing agonist concentrations beyond 10 µM resulted in decreased responses indicating an agonist-dependent antagonism (Fig. 2B). ACh was chosen as the agonist for further experiments, as the activation of nAChRs by ACh is the most biologically relevant interaction for a snake toxin-inhibitor to restore. 10 µM ACh was chosen as the control concentration for further experiments, as it was the lowest concentration that produced the highest level of fluorescence (typically 150-250 RFUs), providing the largest signal window for further experiments, while avoiding an oversaturating concentration of ACh.

**Fig. 2.**
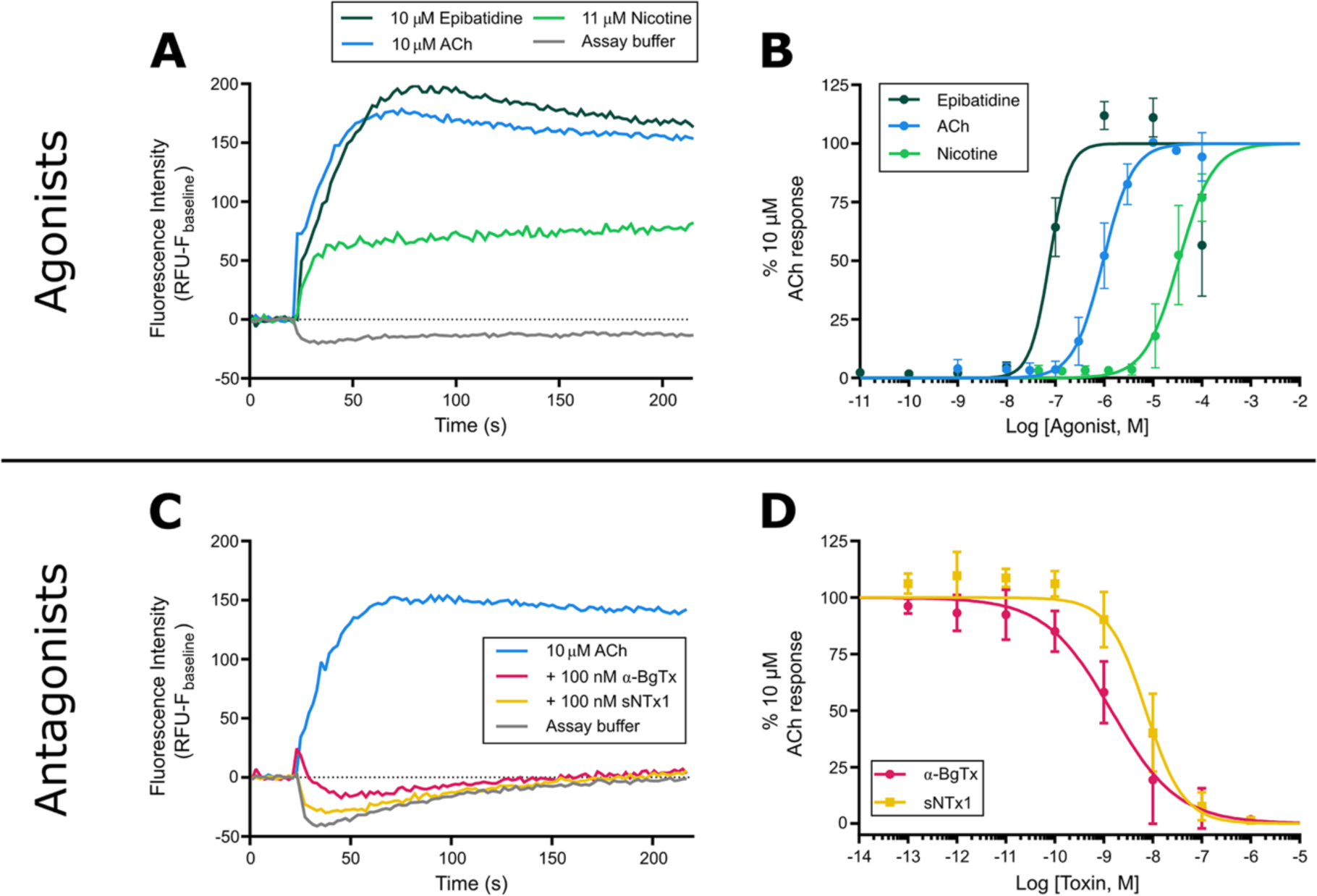
Known nAChR agonists and antagonists show expected action on TE671 nAChR activation when using membrane potential dye. (A) Representative traces showing changes in fluorescence intensity of TE671 cells upon addition of nAChR agonists epibatidine (10 µM, dark green), ACh (10 µM, blue), and nicotine (11 µM, light green), as well as assay buffer only (grey), after 20 s baseline recording. (B) Concentration-response plots showing the changes in the peak fluorescence intensity after addition of serial dilutions of epibatidine (dark green), ACh (blue), and nicotine (light green). (C) Representative traces showing changes in fluorescence intensity of TE671 cells after 15 min pre-incubation with 100 nM of the isolated Lc-α-NTx α-BgTx (red) and the Sc-α-NTx sNTx1 (yellow), followed by addition of 10 µM ACh after 20 s baseline recording. Representative traces of 10 µM ACh control (blue) and assay buffer (grey) are also included. (D) Concentration-inhibition plots showing the inhibition of peak fluorescence intensity of the 10 µM ACh response after the pre-incubation of serial dilutions of α-BgTx (red) and sNTx1 (yellow). Each data point in (B) and (D) represents the mean (±SD) of three independent experiments (n=3).

**Table 1.** The EC_50_ and IC_50_ values of known nAChR agonists, isolated snake venom α-NTxs, and crude snake venoms obtained in this study

| **nAChR modulator** |  |  |
| --- | --- | --- |
| *Agonist* | *EC<sub>50</sub> (μM)* | *95% CI (μM)* |
| Acetylcholine (ACh) | 0.95 | 0.86 – 1.05 |
| Nicotine | 34.00 | 26.60 – 43.40 |
| Epibatidine | 0.08 | 0.03 – 0.23 |
|  |  |  |
| *Isolated venom toxins* | *IC<sub>50</sub> (nM)* | *95% CI (nM)* |
| α-Bungarotoxin (α-BgTx) | 1.42 | 0.83 – 2.44 |
| Short neurotoxin 1 (sNTx1) | 7.23 | 4.83 – 10.80 |
|  |  |  |
| *Crude snake venom* | *IC<sub>50</sub> (μg/mL)* | *95% CI (μg/mL)* |
| *Dendroaspis polylepis* | 0.04 | 0.02 – 0.06 |
| *Dendroaspis j. kaimosae* | 0.10 | 0.04 – 0.29 |
| *Dendroaspis j. jamesoni* | 0.14 | 0.08 – 0.25 |
| *Dendroaspis viridis* | 0.59 | 0.22 – 1.37 |
| *Dendroaspis angusticeps* | 4.49 | 2.24 – 9.02 |
| *Naja subfulva* | 0.03 | 0.02 – 0.05 |
| *Naja haje* | 0.57 | 0.30 – 1.08 |
| *Naja nivea* | 0.30 | 0.23 – 0.38 |

Next, we assessed the ability of the assay to detect nAChR antagonism by α-NTxs. Wang et al. [25] previously demonstrated antagonism by α-BgTx, a Lc-α-NTx isolated from the venom of *B. multicinctus*, to the nicotine response of TE671 cells using membrane potential dye [32]. Consequently, α-BgTx was used along with a commercially available Sc-α-NTx, namely sNTx1 from the venom of *N. pallida*, which was used in previous studies to investigate Sc-α-NTx activity on nAChRs under the name ‘toxin α’ and originally thought to be isolated from venom of *N. nigricollis* [46]. We observed concentration-dependent antagonism of the 10 µM ACh response with both α-NTxs (Fig. 2C and 2D), and α-BgTx was selected as the positive control for measuring the nAChR antagonism of neurotoxic snake venoms in downstream experiments due to its extensive prior characterisation [47].

### 3.2 Fluorescent responses show an acceptable level of plate uniformity for assay use in screening campaigns

With the long-term goal of applying our approach as a novel toxin-inhibitor screening assay, the following controls were selected to assess the uniformity of the assay; i) MAX, a maximal signal produced by 10 µM of the agonist ACh, ii) MIN, a minimal signal produced by the co-application of 10 µM ACh with 30 nM of the antagonist α-BgTx, and iii) MID, a medium signal using the co-application of 10 µM ACh with an IC_50_ concentration of the antagonist α-BgTx (3 nM). Using plates interleaved with these controls (a repeating pattern of three columns occupied by one of each of the controls), inter-day and intra-96 well plate uniformity of assays were performed following a previously described approach [44]. The inter-day assessments validated the reproducibility between different cell populations and passage numbers, whilst the intra-96 well plate experiments revealed no major edge or drift effects, which would invalidate the results when utilising all wells in the plates. Examination of the i) average F_max_-F_baseline_; ii) standard deviations (SD), and iii) coefficient of variations (CV) of the control signals showed clear separation in the three control signals within all plates (Fig. 3). In addition to the low CV and SD, these controls allowed for Z prime (Z’) calculations [45], which are common practice in industrial scale drug screening programmes to determine the distribution of MIN/MAX signals and thus provide confidence that false positive or negative results will not occur. The Z prime of each plate (0.56, 0.62, and 0.57) surpassed the industry-accepted threshold of >0.4, as evidenced by the large signal window and small variance between the MAX and MIN readings.

**Fig. 3.**
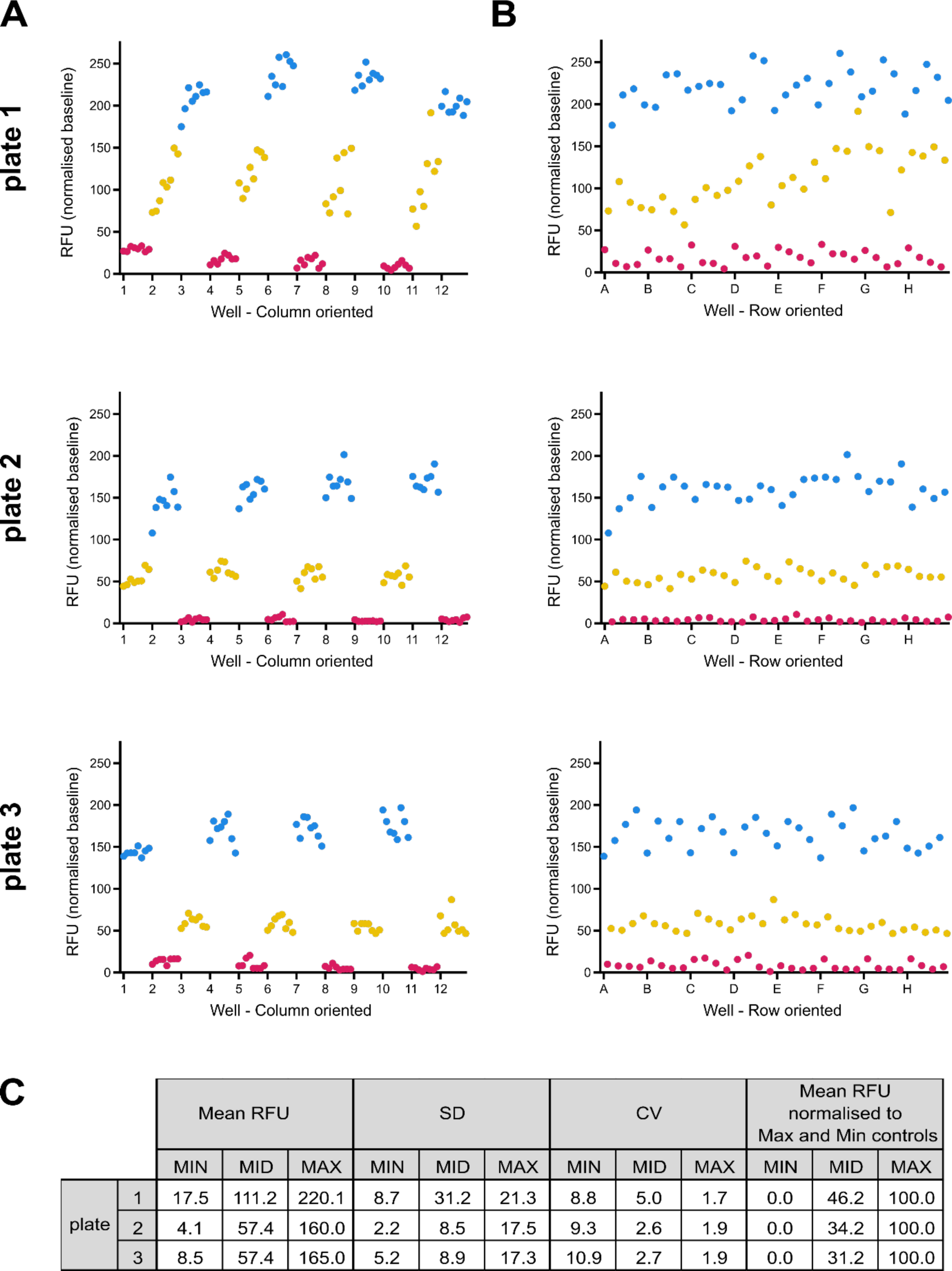
Responses of TE671 nAChRs with membrane potential dye are consistent across a 96-well plate. Scatter plots of the fluorescent response of TE671 cells from wells of a 96-well plate pre-incubated with concentrations of α-BgTx that either give maximal inhibition (30 nM, MIN, red), a middle level of inhibition (3 nM, MID, yellow), or no inhibition (none, MAX, blue), followed by the addition of 10 µM ACh. Each condition was applied to alternating columns of three separate plates (n=3). Each row of plots contains data generated from one plate. Each plate was recorded on a different day with cells of a different passage number and the assigned MIN, MID, and MAX columns were changed on each plate. Each data point is the resulting RFU value after calculating F_max_-F_baseline_ and are plotted by column (A) or by row (B). (C) Summary table of the RFU mean, standard deviation (SD) and coefficient of variance (CV) for each control (MIN, MID, MAX) for each plate. The final column presents the RFU mean once normalised to the MIN and MAX controls.

To assess intra-plate uniformity for each of the three plates, the controls were plotted in spatial order, either by column (Fig. 3A) or row (Fig. 3B). This revealed no consistent drift or edge effects, either across the plate (Fig. 3A – by columns) or down the plate (Fig. 3B – by rows) for all plates. The resulting consistency confirms that responses remain consistent during the read time of the full plate of approximately 40 min where there is a time difference of more than 30 min between the reading of the first and last columns. This validation therefore provides evidence for the use of all wells on the plate, thereby maximising the capacity for multi-plate throughput in a screening campaign. However, inter-plate variation in RFU values after F_max_-F_baseline_ calculation was observed, highlighting the need to normalise readings to the MAX (100% response) and MIN (0% response) control signals to ensure robust cross-plate comparisons. As the entire plate is not read at the same time and responses remain consistent over the recording period, this approach allows the use of a less costly plate reader and is therefore more accessible for many laboratories to implement.

### 3.3 Neurotoxic snake venoms block the ACh response of TE671 cells

Next, we used the developed assay to quantify nAChR antagonism by crude venoms sourced from a variety of medically important African snake species. To this end, we selected eight venoms from cobra (*Naja* spp.) and mamba (*Dendroaspis* spp.) species that are known to contain high abundances of α-NTxs [48–50] and cause systemic neurotoxicity in snakebite victims [11]. All venoms tested showed concentration-dependent antagonism of the TE671 ACh response after pre-incubation with the cells for 15 min (Fig. 4). However, we observed a 100-fold difference in potency across this group of related African elapid snakes (IC_50_s range from 0.03 – 4.49 μg/mL, Table 1, Fig. 4). Venom potency was seemingly not associated with taxonomy, with the rank order of venom IC_50_ from most to least potent being *N. subfulva*, *D. polylepis*, *D. j. kaimosae*, *D. j. jamesoni*, *N. nivea*, *N. haje*, *D. viridis,* and *D. angusticeps* (Fig. 4 and Table 1).

**Fig. 4.**
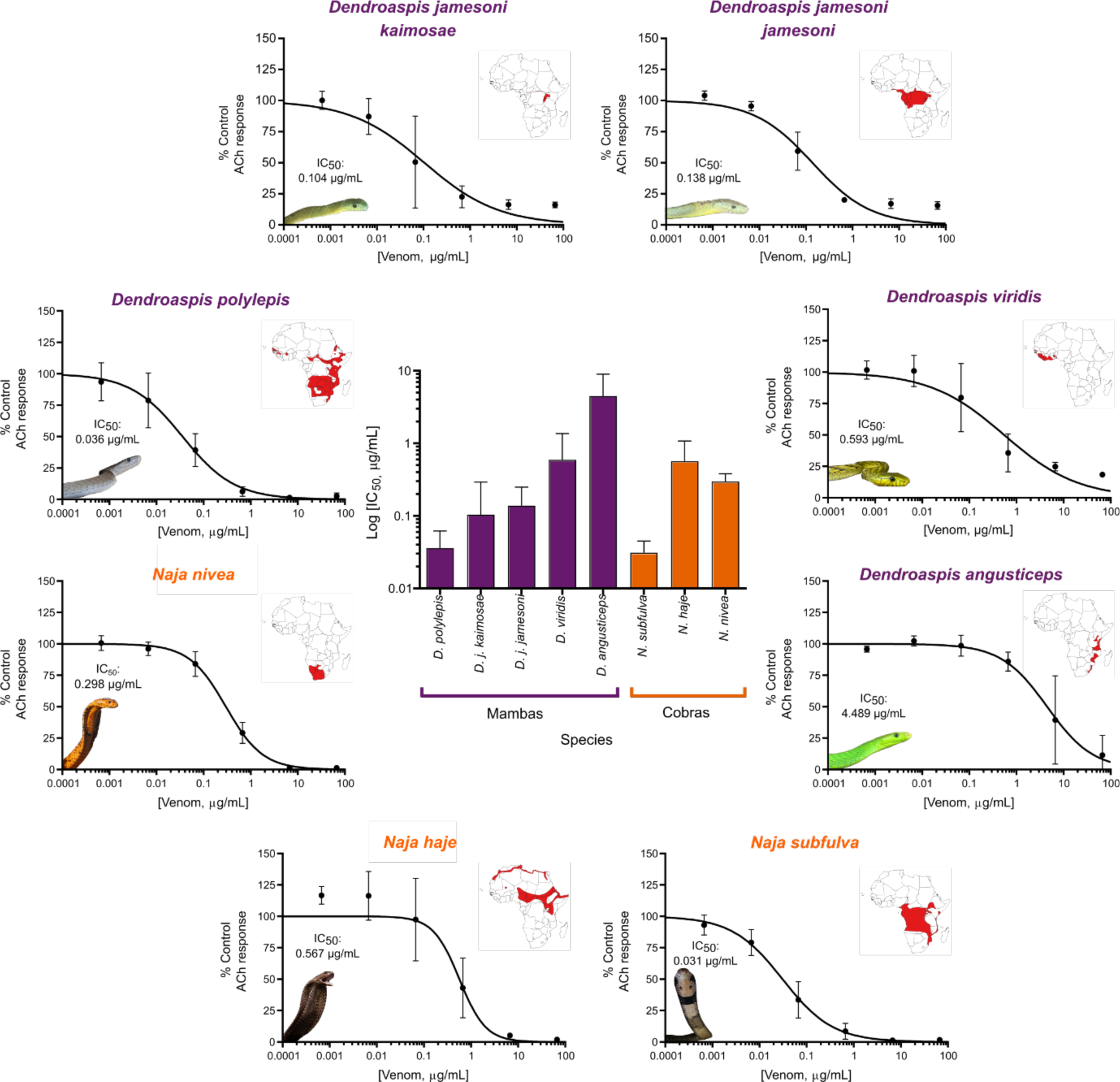
Crude snake venoms show antagonism on nAChRs expressed in TE671 cells. Serial dilutions of crude venom (66.7 – 0.00067 µg/mL) extracted from neurotoxic elapid snake species with geographical distributions covering different regions of the African continent were pre-incubated with TE671 cells for 15 min followed by the addition of 10 µM ACh to create concentration-inhibition plots for each venom (outer ring). Each data point represents the mean (±SD) of three independent experiments (n=3). To the top right of each plot on the outer ring are maps of the African continent highlighting in red the geographical distribution of each species. Maps were generated using QGIS, based on the 2019 International Union for Conservation of Nature Red List of Threatened Species. The central plot compares the IC_50_ values (±95% CI) obtained from mamba (purple) and cobra (orange) venoms and IC_50_ values are displayed above images of snakes inset to the bottom left of each plot.

### 3.4 Different formats of snake toxin-inhibiting molecules rescue the TE671 cell ACh response

In recent years, various molecules have been explored as potential new therapies for snake venom toxins (for a comprehensive overview, see [16,51]). To explore the utility of our assay as a functional screen to detect novel toxin-inhibitory molecules, we selected representatives of these different therapeutic formats (antivenoms, small molecule drugs, nAChR-mimicking proteins, and monoclonal antibodies) and assessed their ability to inhibit the nAChR antagonism stimulated by representative neurotoxic snake venoms (from *N. haje* and *D. polylepis*) and α-NTxs (α-BgTx and sNTx1). In line with the WHO guidelines for preclinical testing of antivenoms [13] and many other *in vitro* and *in vivo* approaches to assess venom inhibition [17,21,28,52], we performed these experiments with an initial pre-incubation step, where inhibitor and venom/toxin were co-incubated at 37 °C for 30 min before assaying, to give the inhibitor maximal opportunity to exhibit neutralisation (Fig. 5). To that end, we also performed these experiments with inhibitory molecules pre-incubated with the lowest venom or toxin concentrations that exhibited maximal nAChR antagonism (6.67 µg/mL for *N. haje* venom, 0.67 µg/mL for *D. polylepis* venom, and 30 nM for α-BgTx and sNTx1). This approach ensured the largest separation between agonist only (100%) and venom/toxin only (0%) signals and that there was not an oversaturating concentration of venom/toxin. In most cases, the inclusion of a 30 min pre-incubation step resulted in a modest increase in the venom/toxin only response (∼10-15% of the agonist only response) (Fig. 5) compared to the response previously observed without incubation (<5%) (i.e., Figs. 2, 3 and 4).

**Fig. 5.**
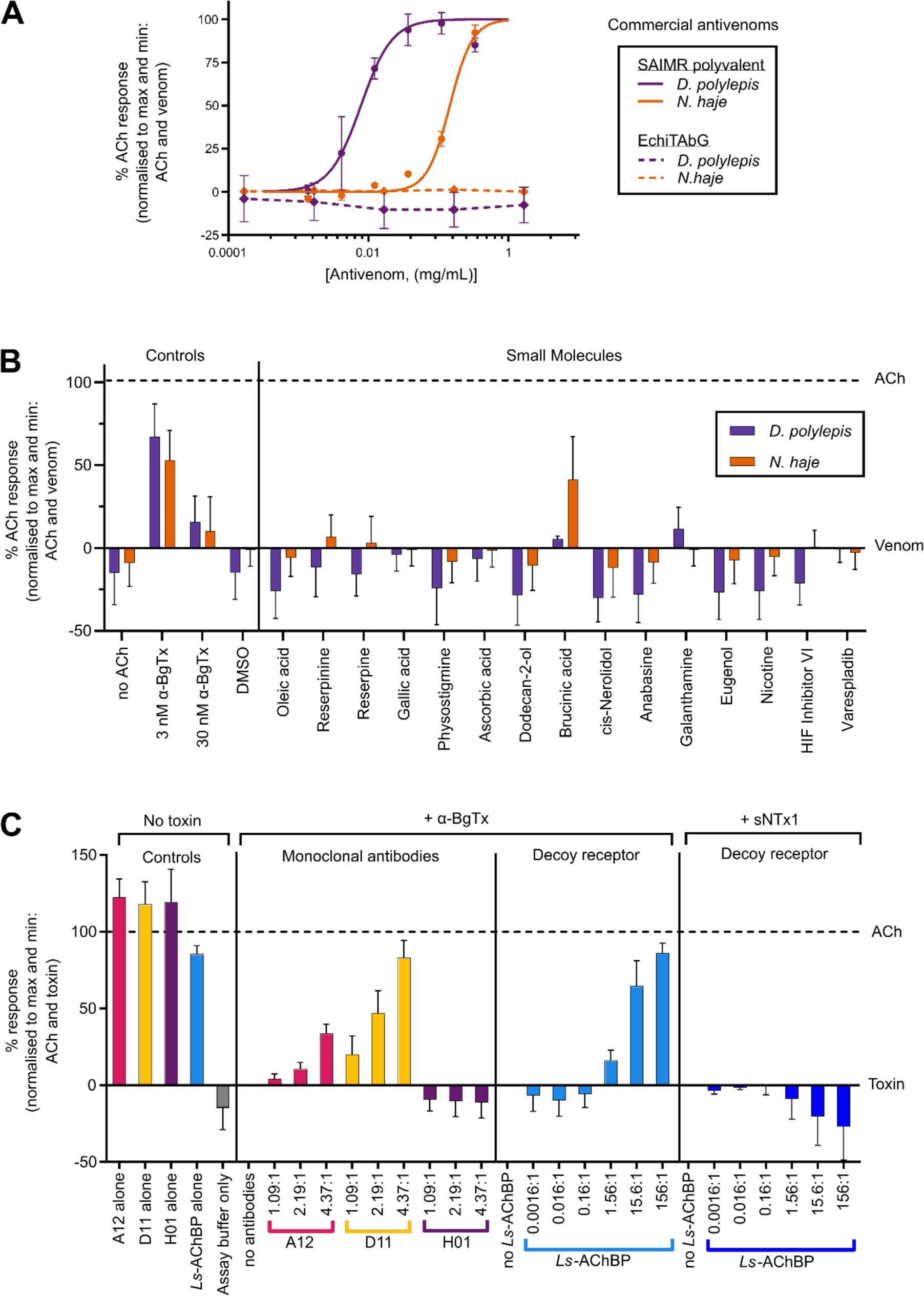
**Inhibition of venom or isolated α-NTxs by commercial antivenom, small molecules, monoclonal antibodies, and decoy receptors**. α-NTx-inhibitors of various formats were co-incubated with concentrations of crude venom (0.67 µg/mL for *D. polylepis* and 6.67 µg/mL for *N. haje*) or isolated α-NTx (30 nM) that gave approximate maximal inhibition prior to application to TE671 cells. All data points represent the mean (±SD) of three independent experiments (n=3) and are normalised to 10 µM ACh (100% signal) and ACh + venom/α-NTx (0% signal) controls with α-NTx-inhibitory activity represented by recovery towards the 100% ACh signal. Venom/α-NTx concentrations were constant at 6.67 and 0.67 µg/mL for *N. haje* and *D. polylepis* respectively. (A) Concentration-response curves showing TE671 ACh response after crude *D. polylepis* (purple) and *N. haje* (orange) venoms were co-incubated with serial dilutions of SAIMR polyvalent antivenom (solid lines, 333.3 – 1.4 µg/mL) and EchiTAbG (dotted lines, 1670.0 – 0.2 µg/mL). Only SAIMR polyvalent antivenom showed α-NTx-inhibiting activity with EC_50_s of 7.8 µg/mL for *D. polylepis* and 146.1 µg/mL for *N. haje*. (B) Screening of a panel of rationally selected small molecules at 100 µM co-incubated with crude *D. polylepis* (purple) and *N. haje* (orange) venom. Each experiment included controls of assay buffer only (no ACh), 3 nM and 30 nM α-BgTx, and 1% DMSO as the drug vehicle control (DMSO). Only brucinic acid (NSC 121865) showed α-NTx-inhibiting activity after incubation with *N. haje* venom, but *D. polylepis* venom was not inhibited. (C) Serial dilutions of the ‘decoy receptor’ *Ls*-AChBP were pre-incubated with either 30 nM α-BgTx or 30 nM sNTx1 at molar ratios ranging from 156:1 – 0.0016:1 and dilutions of the mAbs 2551_01_A12 (A12) and 2554_01_D11 (D11) specific to Lc-α-NTXs, and 367_01_H01 (H01) specific to dendrotoxins were co-incubated with 30 nM α-BgTx at molar ratios of 4.37:1, 2.19:1 and 1.09:1. Inhibition of α-BgTx activity was observed after *Ls*-AChBP was co-incubated with α-BgTx at molar ratios of 156:1 – 1.56:1 but no inhibition of sNTx1 was observed after further dilution. Inhibition of α-BgTx activity was observed with only 2554_01_D11 (restoration to 83.1% of control ACh response with 4.37:1 ratio) and 2551_01_A12 mAbs, with 2554_01_D11 showing a greater level of α-NTx inhibition than 2551_01_A12 (restored to 83.1% of control response as opposed to 33.6% with 4.37:1 ratio). To ensure the α-NTx-inhibitors themselves had no effect on nAChR activation, controls of mAb only (2551_01_A12, 2554_01_D11, and 367_01_H01 alone) and *Ls-*AChBP only at the highest concentrations used for pre-incubation with α-NTxs were also included.

#### 3.4.1 Commercial antivenoms

Commercial polyclonal antibody-based antivenoms are currently the only available specific treatment for snakebite envenoming and are produced by immunising animals with sub-toxic doses of either a single or multiple venoms, resulting in monovalent or polyvalent antivenoms, respectively [13]. We assessed the capability of the assay to detect venom toxin inhibition using SAIMR polyvalent antivenom, which is manufactured using venoms from multiple African cobra and mamba species in the immunisation mixture and, based on prior preclinical research, is known to inhibit venom neurotoxins [53–55]. We used the monovalent antivenom EchiTAbG as a control antivenom, and we did not anticipate observing venom inhibition with this product, as it is specific to the toxins found in the venom of the unrelated, non-neurotoxic, saw-scaled viper, *Echis ocellatus* [56]. Serial dilutions of each antivenom were co-incubated with *D. polylepis* and *N. haje* venom, and responses to ACh addition were compared with responses obtained with venom alone. As anticipated, SAIMR polyvalent antivenom demonstrated concentration-dependent inhibition against the nAChR antagonism caused by both snake venoms, while the non-specific control antivenom EchiTAbG did not show inhibitory activity at any of the concentrations tested (Fig. 5A). Interestingly, SAIMR polyvalent antivenom exhibited greater inhibition against *D. polylepis* venom than *N. haje* based on the lower EC_50_ value, equating to a mass ratio of 1:11.7 (venom:antivenom) against 1. *D. polylepis* and 1:21.9 against *N. haje*, when accounting for differences in venom challenge doses. These findings indicate that more SAIMR polyvalent antivenom is required to neutralise the antagonism of *N. haje* venom, which might be explained by the presence of *D. polylepis*, but not *N. haje* venom in the immunising mixture used to generate this antivenom (instead venoms from related *Naja* spp. are used). Alternatively, the observed differences in neutralising potencies could be due to the considerably higher abundance of α-NTxs in *N. haje* venom [54,57].

#### 3.4.2 Small molecule drug candidates

Next, a panel of small molecules that consisted of either a component from a plant extract that previously demonstrated neutralising activity [40] or that were identified through molecular docking studies of a chemical library with a Lc-α-NTx (α-BgTx or α-cobratoxin from *N. kaouthia*) [41,42] were investigated for their neurotoxin-inhibiting activity (Fig. 5B). Also included in the panel were known nAChR modulators and, as a control, varespladib which is a small molecule inhibitor that exhibits potent inhibition of a different family of toxins found in snake venoms (PLA_2_s), and which is in clinical development [20]. All small molecules were pre-incubated with venoms at a concentration of 100 µM, but only brucinic acid (NSC 121865) exhibited inhibitory activity. Further, inhibitory effects were only observed against *N. haje* venom, where the response was recovered to 41.3% of the control response (Fig. 5B). However, no α-NTx-inhibiting activity was observed for brucinic acid against *D. polylepis* venom.

#### 3.4.3 nAChR-mimicking proteins

AChBPs are soluble proteins found in several mollusc species, and the variant found in *Lymnaea stagnalis* (*Ls*-AChBP) shares features with the human α7 nAChR [58]. A recent study showed that *Ls*-AChBP can bind Lc-α-NTxs from various crude snake venoms, and thus shows potential to act as a decoy molecule that can intercept toxins targeting nAChRs and prevent or delay neurotoxicity [17]. Considering this and previous data showing that Lc-α-NTxs possess a much higher affinity for the α7 nAChR than Sc-α-NTxs [46], we measured the ability of *Ls*-AChBP to inhibit the effects of α-BgTx and sNTx1 in the assay. α-NTxs were pre-incubated with serial dilutions of *Ls-*AChBP, and inhibition of α-BgTx activity was detected (Fig. 5C). Molar ratios were calculated based on an approximate molecular mass of 25 kDa for the *Ls*-AChBP monomer, and the highest molar ratio (α-BgTx:*Ls*-AChBP) of 1:156 restored activity to 86.0% of the ACh control. The lowest ratio to exhibit any restoration was 1:1.56 (16.2% of ACh control). As anticipated, we observed no inhibition of the antagonism of sNTx1 at any of the tested *Ls*-AChBP concentrations (Fig. 5C).

#### 3.4.4 Monoclonal antibodies

A recent study identified the mAbs 2551_01_A12 and 2554_01_D11 as effective inhibitors of several Lc-α-NTxs, including α-BgTx, using automated patch-clamp electrophysiology and murine *in vivo* experimentation [21]. Consequently, we used our assay to explore whether the α-NTx inhibition of these mAbs could also be detected in this assay, using α-BgTx as our model (Fig. 5C). Pre-incubation of solutions containing 1:1.09, 1:2.19 and 1:4.37 molar ratios (α-BgTx:mAb), calculated based on using 150 kDa as an approximate molecular mass for each IgG1 mAb, were tested, alongside a negative control mAb (367_01_H01) directed against dendrotoxins from *D. polylepis* venom [59]. Concentration-dependent inhibition of α-BgTx activity was observed with both antibodies directed against Lc-α-NTxs (2551_01_A12 and 2554_01_D11), in line with previous electrophysiological findings [21]. The mAb 2554_01_D11 was able to restore nAChR activity to a higher percentage of ACh control (83.1%) than 2551_01_A12 (33.6%) at the highest dose tested. As anticipated, the control anti-dendrotoxin mAb (367_01_H01) exhibited no inhibition of α-BgTx antagonism of the nAChR, even at the highest concentrations tested.

## 4. Discussion

In this study, we developed a cell-based assay to investigate venom toxin activity on muscle-type nAChR activation and explored its capability to detect inhibition by various toxin-inhibiting molecules. For validation, we first quantified the effects of known nAChR agonists and antagonists to ensure that their observed effects were consistent with other validated experimental techniques, and that any modifications made to previously published approaches [31,32] did not affect the assay (Fig. 2). The time to peak and decay of fluorescent responses during the recording time (Fig. 2A) were consistent to those observed in other studies employing the same experimental approach [60], though differences were observed when comparing outcomes with electrophysiology approaches. Responses of muscle-type nAChRs typically reach a peak and return to baseline within a few seconds in electrophysiology experiments [61], while the responses observed in this assay do not return to baseline after 214 s of recording (Fig. 2A and 2C). This prolonged response could perhaps be due to a positive allosteric modulatory effect of the dye or the activation of natively expressed voltage-gated ion channels [43] after nAChR activation. Irrespectively, the profile of responses and the sensitivity of the assay to traditional nAChR modulators confirms the utility of this approach for measuring nAChR activation, rather than other physiological properties of the channel. The EC_50_s for agonists (Table 1) also differ from those obtained with electrophysiology approaches [33,62–64], but remain consistent with those obtained in previous studies using the same experimental approach [31,37]. Antagonism by α-BgTx was confirmed as in previous studies (Fig. 2C and 2D) [32,65], and sNTx1 also exhibited antagonism (Fig. 2C and 2D), which was anticipated given that binding of this α-NTx to the muscle-type nAChR has previously been demonstrated [46]. Collectively, these data provide confidence that the developed assay is informative for assessing nAChR agonism and antagonism.

All snake venoms tested in this study (from *Naja* and *Dendroaspis* spp.) showed evidence of antagonism on the muscle-type nAChR (Fig. 4). These findings were anticipated, since: (i) systemic envenoming by these species result in neurotoxic clinical manifestations in snakebite patients [11,12], (ii) α-NTxs have previously been identified in various mamba and cobra venoms [48–50,54,55], and (iii) venoms from *N. haje* and *D. polylepis* have previously been demonstrated to exhibit nAChR antagonism in functional assays [32,66]. Since there was an almost 100-fold difference in potency of the crude neurotoxic venoms investigated in this study, with no obvious correlation with taxonomy, investigation of additional elapid venoms from diverse genera could be particularly revealing to unravel the evolutionary basis of these considerable differences in venom potency. Given the medical importance of α-NTxs, the assay described here could be readily used in conjunction with venom fractionation/purification and identification approaches [55,67] to identify the key α-NTxs responsible for nAChR-mediated neurotoxicity. Such ‘toxicovenomic’ profiling is important, as each snake venom can potentially contain multiple α-NTxs, and these likely differ in both potency and abundance, as well as they may vary both intra- and inter-specifically [49,54,68]. The identification of such toxins is important for the rational selection of targets for novel therapeutics (antivenoms and toxin-inhibitory molecules), and/or to either supplement or use as alternatives to, whole venoms as immunogens for antivenom production [18,69]. Additionally, given that toxins outside of the 3FTx family have also been demonstrated to exert nAChR antagonism [70], this approach may also prove useful for identifying novel venom neurotoxins.

The assay was further demonstrated to be compatible with the detection of ability of various therapeutic candidate molecules to inhibit the antagonism of venom neurotoxins on the nAChR (Fig. 5). In several cases, pre-incubation with venom or α-NTxs resulted in a restoration to >80% of the control response, demonstrating clear inhibition (e.g., SAIMR polyvalent antivenom, *Ls*-AChBP, and mAb 2554_01_D11). Given the demonstrated acceptable level of uniformity across a 96-well plate (Fig. 3), there is clear potential to use this assay as a screening platform for the identification of novel toxin-inhibiting molecules against venom nAChR-antagonists. For example, in an approach analogous to that proposed elsewhere for other venom toxins [71], this assay could be implemented as a primary drug screening assay to identify α-NTx-inhibiting molecules present in compound libraries consisting of drugs that are already approved or in development for other indications. This ‘drug repurposing’ approach is particularly attractive for snakebite envenoming, as ensuing hits have often entered at least early-stage clinical trials for other indications, resulting in potentially shorter development timelines compared with the development of new chemical entities, and therefore potentially lower development costs [71]. Similarly, this assay could be used for aiding the discovery and optimisation of cross-reactive mAbs directed against α-NTxs and other 3FTxs. Such approaches currently rely on binding assays for screening [72], typically followed by complex and expensive bioassays (e.g., patch-clamp electrophysiology or *in vivo* preclinical studies) to assess α-NTx inhibition [21]. The same principles apply to the development of receptor-mimicking peptides/proteins based around AChBP scaffolds. Recent insights into the properties of Sc-α-NTx binding to muscle-type nAChRs [61] should aid the future protein engineering of AChBP derivatives and receptor-mimicking peptides designed to simultaneously capture both Lc-α-NTxs and Sc-α-NTxs. Such molecules could be readily screened in this assay for their generic α-NTx-inhibiting activity as an initial readout to inform downstream structural optimisation and lead candidate selection.

The use of the human muscle-type nAChR in this assay is a particular strength, as questions have been raised about the appropriateness of rodent models for assessing the activity of α-NTxs due to Sc-α-NTxs exhibiting enhanced potency on rodent nAChRs [73]. Investigating human nAChRs in TE671 cells can help ensure that only toxins relevant to causing neurotoxicity in snakebite victims are being studied. However, certain α-NTxs can exhibit enhanced potency for foetal (γ subunit-containing) muscle-type nAChRs (subtype expressed in TE671 cells) over the adult (ε subunit-containing) type, as previously observed with the Sc-α-NTx ‘NmmI’ from *N. mossambica* [74]. The CN21 cell line, used in a similar cell-based fluorescence assay to investigate chemicals to counteract organophosphate poisoning [75], expresses the adult muscle-type nAChR and could be employed in place of TE671 cells to distinguish α-NTxs selective for foetal muscle-type nAChRs. Another future expansion of this assay would be to employ a more challenging model of envenoming. After identifying promising α-NTx-inhibiting candidates in pre-incubation experiments, these candidates could then be assessed using a model where venom/toxin is applied simultaneously or before the toxin-inhibitor. This is relevant, because a major hurdle for α-NTx treatments to overcome is the long dissociation time of Lc-α-NTxs once bound to the nAChR [68]. Studies using chick biventer cervicis nerve-muscle preparations and commercial antivenoms have employed an analogous approach and showed the ability of antivenoms to reverse α-NTx dependent inhibition of nAChRs when applied after treatment with Asian cobra venoms [76] and *Oxyuranus scutellatus* venom [77]. Adaptation of our assay in a similar manner could allow for further discrimination between inhibitors that promote toxin dissociation from the nAChR compared with those that need to intercept α-NTxs before they bind.

The herein described approach of measuring TE671 cell muscle-type nAChR activation using membrane potential dye has enabled the assessment of nAChR antagonism by crude elapid snake venoms and isolated Lc-α-NTx and Sc-α-NTxs. As both classes of α-NTxs exhibited dose-dependent antagonism, the assay provides a robust platform to investigate toxicity mediated by α-NTxs from the venoms of snake species found across different geographical regions. This assay could also find wider utility for studying nAChR modulators, whether from natural (e.g., other animal venoms or toxins) or chemical sources. In addition, we demonstrated the utility of the assay for identifying α-NTx-inhibitory molecules and highlight its compatibility with four major categories of snakebite therapeutics currently being explored. We therefore hope that this assay will be a useful addition to the experimental toolbox to identify new therapeutics against key neurotoxins from snake venoms, and that it will help deliver new toxin-inhibitors that can mitigate the many life-threatening snakebite envenomings that occur worldwide each year.

## Acknowledgments

We thank David Richards for early discussions on the assay concept, Ian Mellor (University of Nottingham, UK) for the gift of TE671 cells, and IONTAS (Cambridge, UK) for facilitating access to the monoclonal antibodies. We also thank Jory van Thiel for the creation of species distribution maps presented in Fig. 4, Paul Rowley and Edouard Crittenden (Liverpool School of Tropical Medicine, UK) for expert maintenance of venomous snakes and provision of venom samples, as well as use of photographs presented in Fig, 4, alongside those generously provided by Taline Kazandjian, Steven Hall, Simon Townsley, and Wolfgang Wüster. This work was supported by the Wellcome Trust funded project grants 221710/Z/20/Z (CU, JK and NRC) and 221712/Z/20/Z (NRC and JK) and KU Leuven grant C32/16/035 (CU). MN is a recipient of a FWO postdoctoral fellowship 12X2722N. This research was funded in part by the Wellcome Trust. For the purpose of open access, the authors have applied a CC BY public copyright licence to any Author Accepted Manuscript version arising from this submission.

